# Exploration in the presence of mother in typically and non-typically developing pre-walking human infants

**DOI:** 10.1101/350736

**Authors:** T. Frostig, H. Alonim, G. Scheingesicht, Y. Benjamini, I. Golani

## Abstract

Using an arsenal of tools previously developed for the study of origin-related exploration in animals, we compared exploration of human pre-walking Typically-Developing (TD) and Non-Typically Developing (NTD) infants in the presence of mother. The NTD infants had been referred to a center for the treatment of autism by pediatric neurologists and expert clinicians. Using computational analysis we document in TD infants a phylogenetic ancient behavior: origin-related exploration. Strikingly, while the TD infants exhibited excursions in reference to mother and deep engagement with mother when visiting her, the NTD infants tended to avoid mother’s place, performing few if any excursions, and exhibiting shallow engagement with mother. Given the pervasiveness of origin-related exploration in invertebrates, vertebrates, and primates, we now face a challenge to find an animal model that will exhibit active exploration while ignoring or suppressing the return to the origin, be it a mother or any other safe haven.

## Main

In this study we sought to establish the generative rules that shape the path traced by human pre-walking infants in moment-to-moment exploration of a novel environment, under the protective attention of the infant’s relatively stationary mother. Our perspective comprised three traditionally distinct fields of study: that of animal exploratory behavior, that of primate- and that of human mother-infant interaction.

A conspicuous spatial regularity in the exploratory behavior of many organisms is that of a reference place in relation to which they explore the environment. In the wild, many animal species have a home site to which they return regularly after exploring their home range or territory, be they, for example, wolves (Fritts & Mech 1981), small mammals (Brown 1966), ants (Martin & Rudiger 1988) bumble bees (Woodgate et al. 2016), or millipedes (Hoffmann 1984). In behavioral neuroscience experiments, rats have been shown to explore the experimental arena from a reference place, from which they perform excursions into the environment (Eilam & Golani 1989). The high accumulation of time spent across a large number of visits also characterizes the reference place of, for example, mice (Fonio et al. 2009), zebra fish (Stewart et al. 2010;Stewart et al. 2011), and infant rats (Loewen et al. 2005). This reference place, often termed a "home base", exerts its influence on the organism’s behavior across the entire exploratory basin. Visits to the home base partition the path into separate excursions in the environment. The latter are further partitioned into progression segments and staying-in-place (lingering) episodes (Drai & Golani 2001). In moment-to-moment behavior in a novel environment the excursions grow in extent (Benjamini et al. 2011) and differentiate from simple excursions to complex ones (Benjamini et al. 2011).

The performance of exploratory excursions has also been reported in the wild in infant primates – rhesus monkeys (Berman 1980), baboons (Altmann & Samuels 1992), and chimpanzees (van de Rijt-Plooij & Plooij 1987). As the infants of these primates develop, they perform increasingly longer excursions from mother into the environment and back to mother. While the mother is often on the move during the performance of such excursions, the excursions nevertheless involve both exploration and active management of distance in reference to an origin or base by the infant, be it a mobile or stationary mother.

Origin-related exploration has also been described in human infants: reinforcing Bowlby’s attachment theory with ethological data, Ainsworth observed that human infants "tend to explore on their own initiative an unfamiliar room in their mother’s presence, and in doing so use mother as a secure base" (Ainsworth & Bowlby 1991;Ainsworth 1969). Similarly, Mahler’s psychoanalytic separation-individuation theory on the psychological development of the human infant is largely supported by observations of the infants’ "incessant wandering away" and "checking back to mother", using mother as reference and "home base" (Mahler et al. 1975). Human infant performance of increasingly longer exploratory excursions from a stationary mother has also been reported previously (Rheingold & Eckerman 1970), and in one study also plotted (Vitelson 2005).

The current report presents the kinematics on a moment-by-moment actual-genesis scale, enabling a computational comparison across taxonomic groups of the rules that shape an organism’s path. The study is complementary to the majority of infant mother-related exploration studies, which consist of verbal reports of behavior, or reports of inferred intrapsychic processes, such as symbiosis (Mahler et al. 1975), separateness^20^, identity (Stern 2009), self, object relationships, or separation anxiety (Ainsworth 1969), on a day-by-day or month-by-month developmental scale (Jones 1972;Rheingold & Eckerman 1970). The current study sought to formulate some of the rules that shape mother-related exploration in pre-walking human infants, using a description that discloses the infant’s engagement with the environment surrounding it. To obtain a wider perspective on the behavior of Typically-Developing (TD) infants, we compared the behavior of six TD infants to that of six Non-Typically Developing (NTD) infants referred to the Mifne Center for the assessment and Treatment of Autism (Alonim 2004) - three infants were referred for developmental assessment by expert clinicians and three were referred for treatment by pediatric neurologists. All the NTD infants had primary assessment due to parental concerns regarding their infants’ development.

We perceived a potential byproduct of this comparison as highlighting those behavioral symptoms that could later be looked for in prospective animal models of this subgroup of NTD infants.

We found that origin-related exploration was weak or even absent in the NTD infants. This finding is perhaps not unexpected from the perspective of attachment theory; it is, however, surprising from a phylogenetic perspective, given the pervasiveness of origin- and mother-related exploration across primates, vertebrates, and even arthropods.

Our results provide a comparative phylogenetic perspective, a glance into a conserved, relatively universal structure preceding specific functions it fulfills in human infants, and a glimpse into the distinct operational worlds of the TD and a sub group of NTD infants, and into the ways in which they attend to the world and come to grips with it. We thus start with animals, proceed to human infants and aim at returning to animals

### Dwell time distribution across the room

This study was conducted on pre-walking infants at the stage of stable crawling. The infants were recruited with their respective mothers to participate in a video-taped half-hour session conducted in a medium-sized room. In each trial the mother entered the room carrying her infant, sat on a mattress near the wall, and then seated the infant or let it slide down next to her. The mother was requested to remain seated and allow the infant to act freely for a 30-minutes session. Two video cameras were used across the session: one capturing a view of the whole room including the infant and the mother (fig. 1); and another zooming in on the infant and following it. The behavior of the infant and the mother was video-taped and then tracked for half an hour, a period that had been found to be appropriate in our preliminary study (see *Methods*).

**Figure 1.**
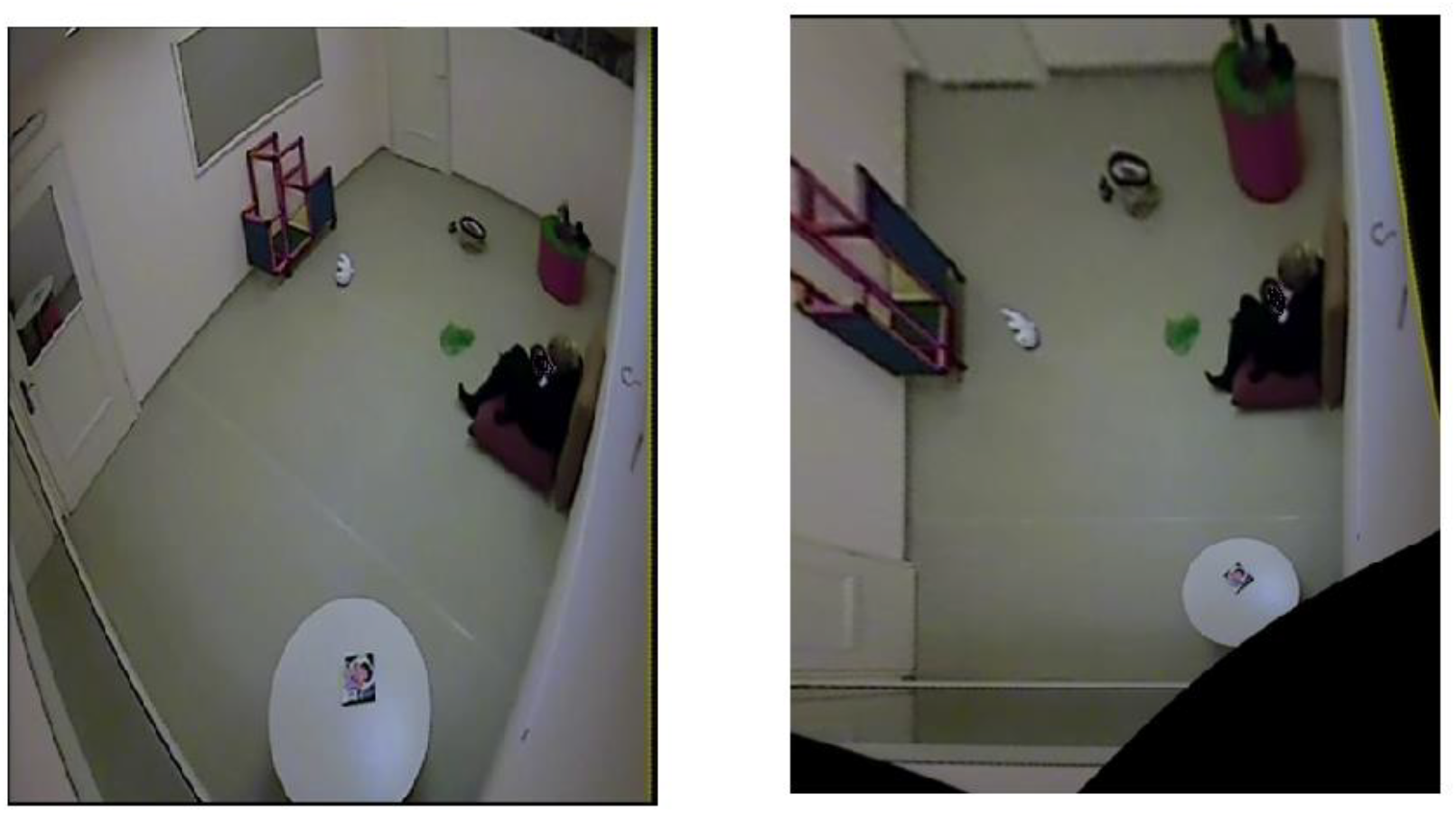
The Mifne Treatment room image, before (left) and after transformations (right). Note furniture, toys, mother’s location (here with infant sitting on her lap), on the right side near the wall), and two doors (on the left – exit doorway and on top door leading to bathroom).

Figures 2a, b present for both the TD and NTD infants the smoothed cumulative dwell-time spent in different locations across the observation room and the number of visits paid by them to those neighborhood locations exhibiting peak dwell times (see *Methods*). Dwell-time is represented by colored contour lines forming a topographical map. A comparison of, for example, the TD infant Alon with the NTD infant Tom (all names provided in this study are fictitious), reveals that, in both, dwell time displays a patchy distribution across the room. However, whereas for Alon there was a single peak, located near mother, which stood out in terms of dwell-time (signified by a yellow center), for the NTD infant Tom there were two peaks of relatively the same dwell-time and both were located away from mother. Alon spent time over the whole room whereas Tom adhered to the upper half, hardly lingering in the lower half of the room. Similar differences were revealed for most of the infants in the respective groups: all the TDs exhibited a single preferred place in terms of dwell-time, whereas the NTDs (except for two: Yuri and Dean) exhibited more than one peak dwell-time place. In all the TDs except for one (Alexey, who also exhibited several NTD features; see Method’s section), the peak dwell-time place was located by mother, whereas in all NTDs it was located away from mother.

**Figure 2.**
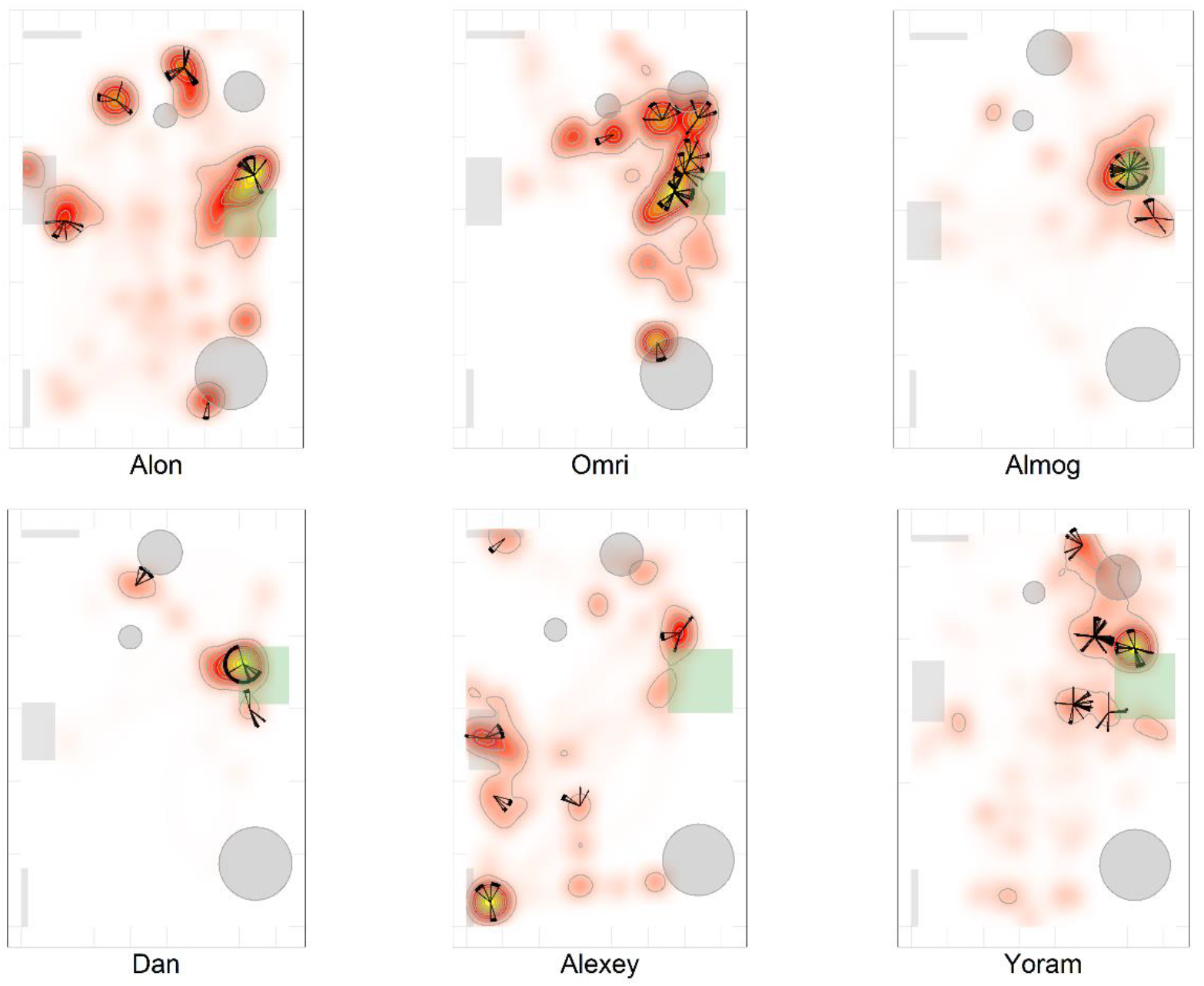
The TD infants’ most preferred place was located near mother and they visited it 175 regularly paying it the highest number of visits. The figure presents the smoothed cumulative 176 dwell-time spent in different locations across the observation room, and the number of visits 177 paid by the infants to neighborhood locations exhibiting peak dwell-times. Number of visits is 178 obtained using the two concentric circles method (*r*_*in*_ = 30*cm*, *r*_*out*_=50*cm*, see Methods). 179 Dwell-time is represented by colored contour lines forming a topographical map. The contour 180 lines are spaced at the quantiles (0.1, 0.2, 0.3, …) of the smoothed cumulative dwell-time (see 181 Methods). Note that in these heat maps the color is proportional to the dwell time of each 182 specific infant.

**Figure 2b.**
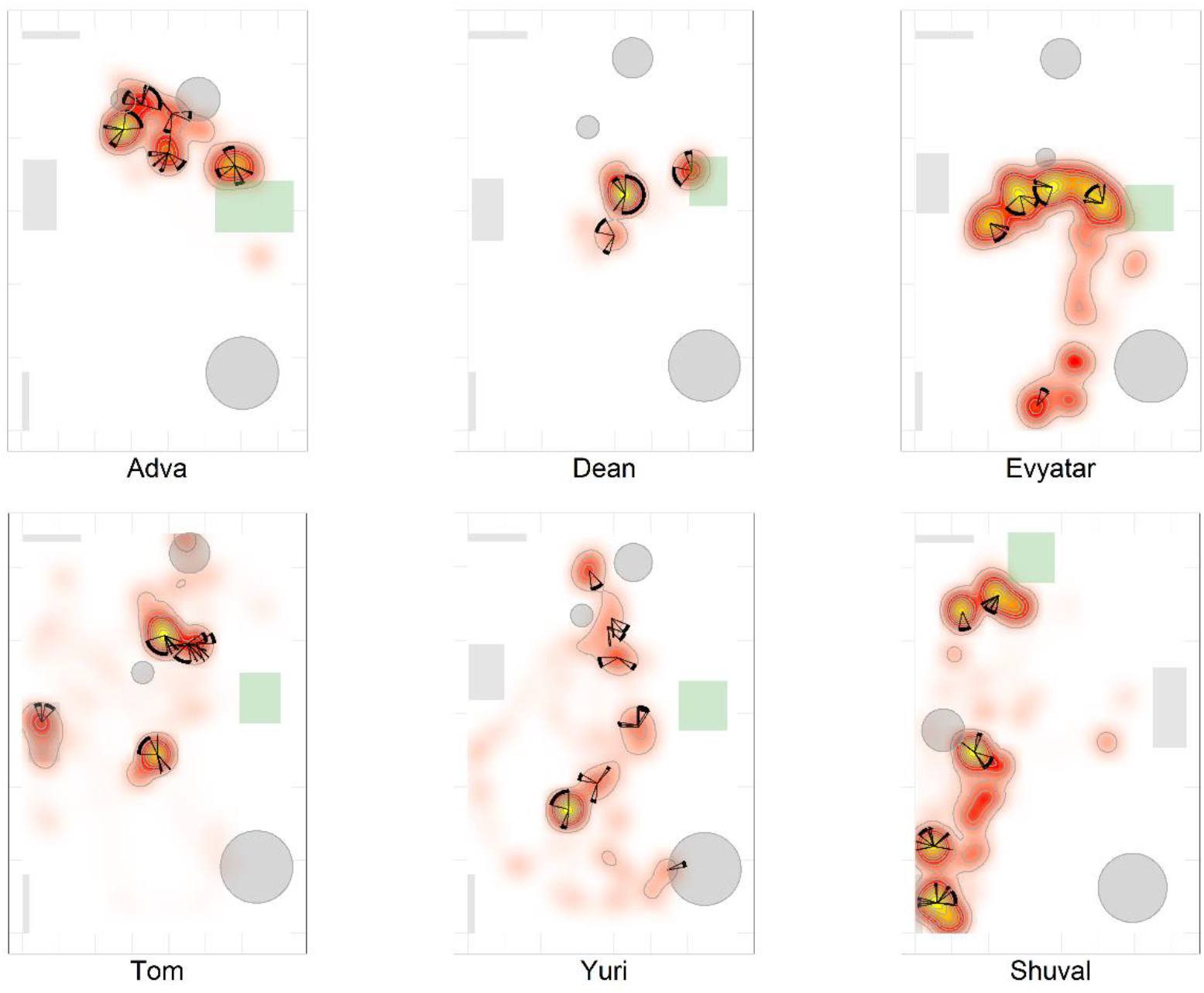
The NTD infants’ most preferred places were not located near mother and the 186 infants did not visit these places regularly, nor necessarily paying to them the highest number 187 of visits. For further explanation see Figure 2a.

Whereas the TD infants tended to establish preferred places also near furniture (Alon by the cabinet, table, and basket; Omri by the cylinder and table; Yoram and Dan by the cylinder; and Alexey by the cabinet and the exit door), the NTDs often established places away from both walls and furniture. Five of the NTD infants (Evyatar, Tom, Dean, Yuri, and Shuval) established peak dwell-time places in the open space facing their mother, but without returning to her. One NTD infant (Shuval) and one TD (Alexey), established peak dwell times *vis-à-vis* the door leading out of the room, perhaps disclosing an intention to leave the room.

While the TDs tended to cover the whole room (including Dan, whose extended staying with mother dwarfed other staying-in-place episodes across the room; see also animations), the NTDs tended to adhere to only part of the room (See *Endpoint Summaries* - % room covered): Adva to about a third of the room, completely ignoring the other parts; Dean to the center part of the room vis-à-vis mother, ignoring all other parts; Evyatar first in the lower part of the room and then spending all his peak dwell-times in a horseshoe-shaped strip vis-a-vis mother, ignoring the rest of the room; and Shuval ignored the right half of the room. Of the NTDs, only Yuri covered most of the room and this appeared to be due to his chasing a large rolling ball for most of the session.

### Visit distribution across high dwell-time places

Figures 2a, b provide a spatio-temporal summary of the process by which dwell time was *allocated* across the session to the main places. The stars on top of the peak places represent the number, order, and duration of the visits to the respective peak dwell-time places. Starting at 12 o’clock, the circumference represents the session’s half-hour duration. Proceeding clockwise, each wedge presents a visit to this location, with its start and end times given by the orientation, and its duration thus presented by the arc’s length. For very short visits the wedge appears as a single spoke. Almog (TD), for example, visited the most preferred place, located by his mother, 14 times, thus paying the highest number of visits to the place near mother and accumulating time near mother in a piecemeal manner, whereas Dean (NTD) visited his most preferred place only 3 times, ignoring that place for a third of the session and visiting mother only twice. These differences in visit management apply between most of the two groups of infants: all TDs except for Alexey (whose peak visiting place was the exit doorway) most visited place was near mother for the highest number of times; whereas all the NTDs’ most visited places were located away from mother and each was typically ignored for substantial parts of the session. In short, the NTD infants did not show sustained attention to any place, or object as expressed in spread-across-the-session visits. In both groups visits to most places (except the TD infant mothers’ places) were sparse and not evenly distributed. Both Alexey (TD, but see *Methods*) and Shuval (NTD) paid the highest number of visits to the exit doorway.

It should be noted that the number of visits to peak dwell-time places does not exhaust the number of visits to mother, and therefore does not disclose the full number of excursions performed from the mother; these visits merely refer to one or at most two places in her vicinity. The infants might, and indeed did, visit mother’s vicinity from other, less visited directions, not necessarily belonging to the peak dwell time places.

The stable preference of the TD infants for the place located near the mother and the absence of a preferred place near the NTD infants’ mother (fig. 2a, b) marks the significance of the TD mothers’ location for the behavior of the TD infants, thus requiring an examination of all the infants’ path-sessions in explicit reference to mother’s location (See *Endpoint Summaries* - # of excursions and % time spent near mother).

### The itinerary, duration, and extent of engagement of the infants’ visits to mother

We plotted in the correct order, across the whole session, the relative time of start, the relative time of end, the duration in session percentages, and the extent of engagement by the infant when in mother’s proximity, noting whether the infant merely approached mother during a specific visit; Whether it touched her; and Whether it climbed on her (see *Methods*). As shown, Alon (TD, fig. 3a) visited mother ca. 12 times, sometimes only approaching her and at other times climbing on her; whereas Tom (NTD, fig. 3b) spent the first few seconds at the very start of the session near mother and then later slightly approached her three times. More generally, five of the TD infants (Alon, Omri, Yoram, Almog, and Dan) came to close, extended, and persistent grips with mother, whereas all the NTD infants either avoided mother’s proximity (Tom and Yuri), or avoided visiting her for extended parts of the session (Dean, Evyatar and Shuval). Adva, an NTD infant, did visit mother, but only four times. Note that in 5 TD (Alon, Omri, Almog, Dan, and Yoram) and 2 NTD (Adva and Dean) infants the last extended visit involved, using Mahler’s description (Mahler et al. 1975), "melting into mother". The difference in the extent of engagement with mother is also evident in the plots presented in figure S1 of both the average percent time spent, and the number of visits paid at distances starting from zero centimeters from the center of her location (fig. S1).

**Figure 3.**
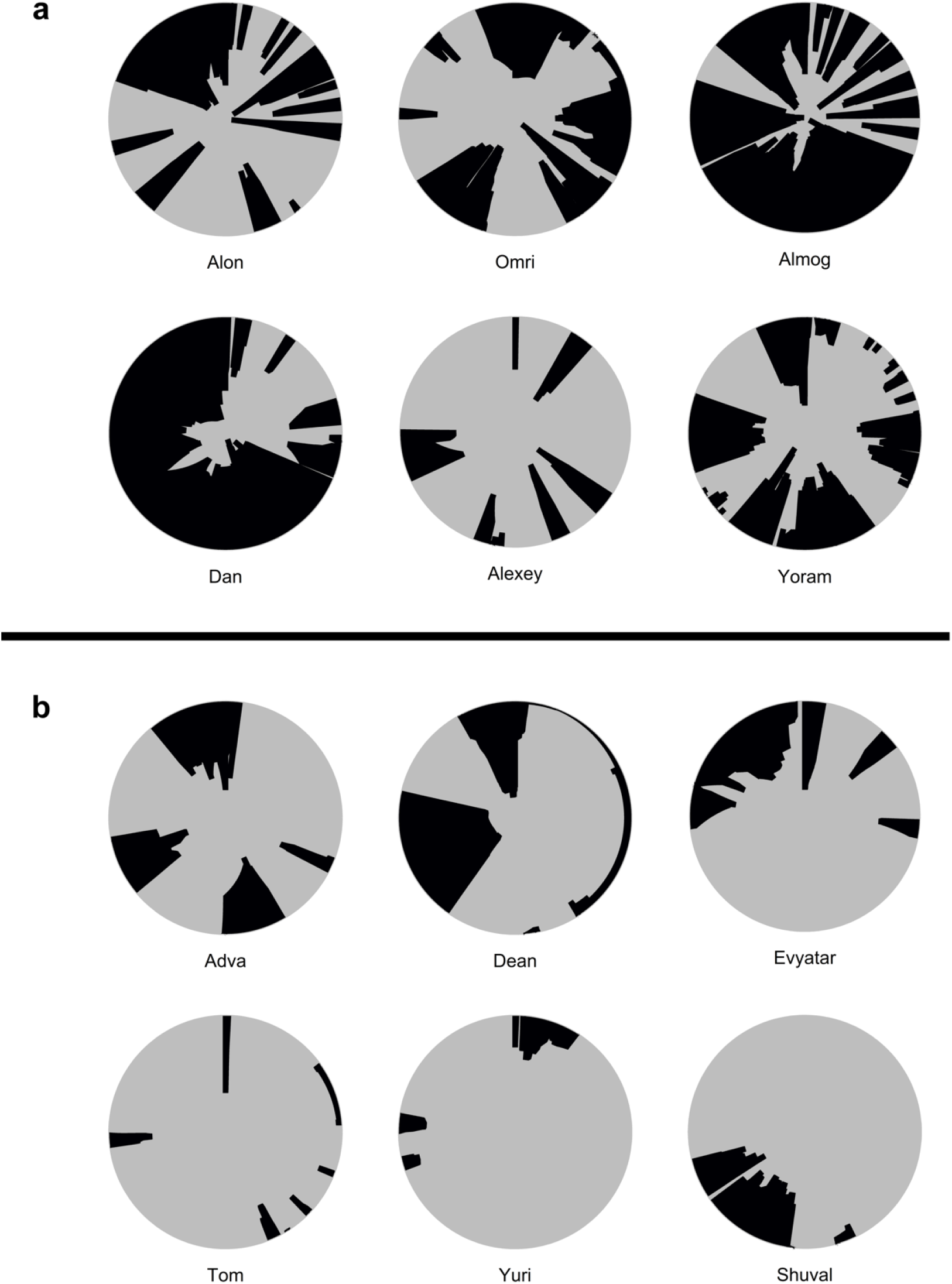
The TD infants (a) visit mother frequently and persistently, engaging her deeply 215 and for long durations, whereas the NTD infants (b) visit mother rarely if at all; and when 216 visiting they typically merely approach and do not touch mother. The engagement plots exhibit 217 the timing, duration, and extent of the infant’s being in gear with mother. The concentric circles 218 are centered on mother’s location, spanning a radius of hundred and twenty cm around that 219 center. Starting at twelve o’clock and proceeding clockwise for the session’s duration, the arc 220 traced on the circle’s circumference and the colored section of the circle designate the times of 221 start, end, and duration, and the extent of engagement with mother’s proximity.

### The itinerary, duration, and extent of engagement of the infants’ visits to furniture in the room

Figure 4a, b presents plot summaries of the extent of engagement of each of the infants with mother, with the four items of furniture in the room, and with the two doorways leading out of the room: how each of the infants comes to grips with structures in the environment, and to what extent they come to grips with mother.

**Figure 4.**
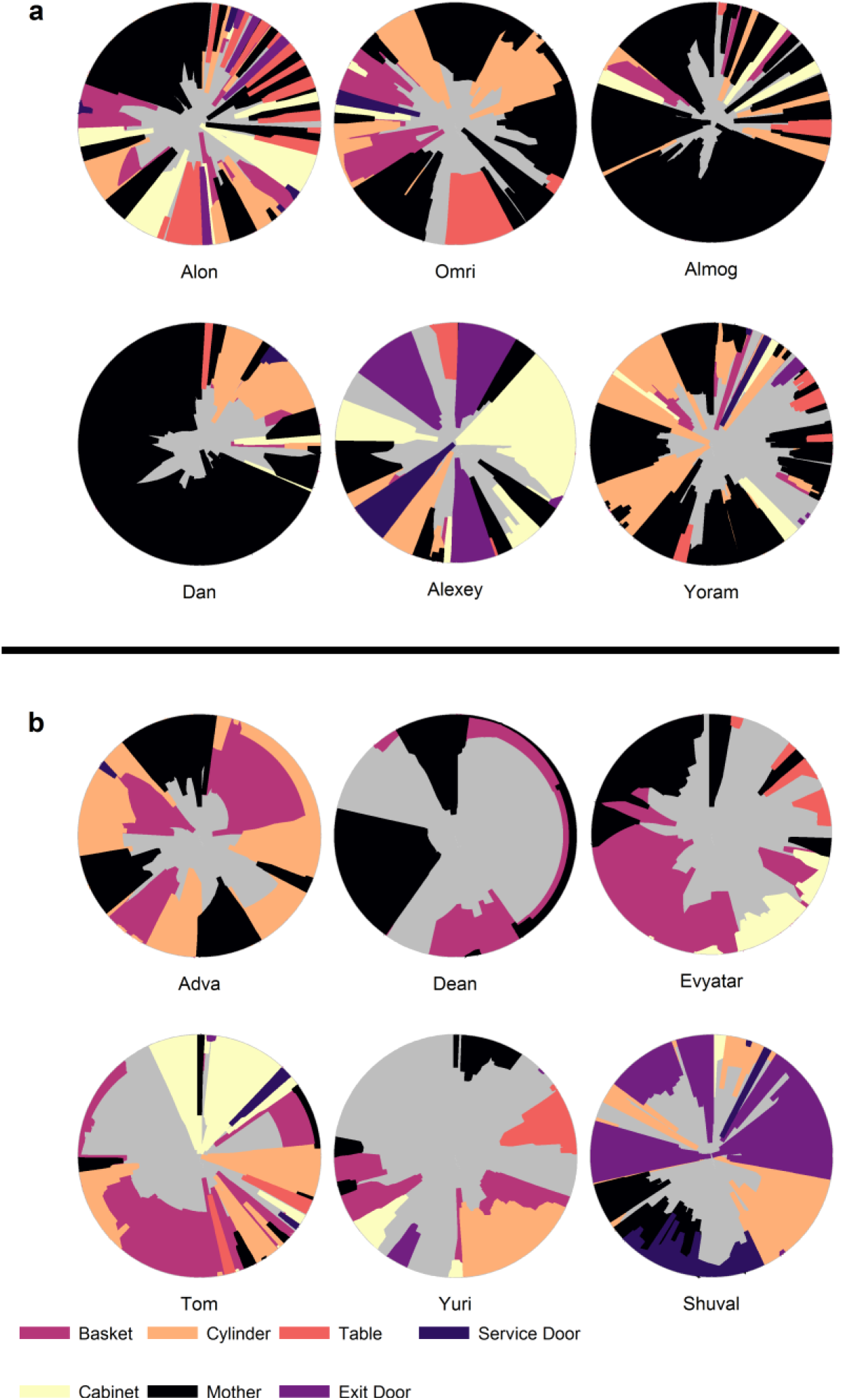
The TD infants visited mother and the furniture items frequently and persistently, 231 invading their respective places deeply and for long durations. b The NTD infants’ visits to 232 mother and to the furniture were infrequent and shallow. For explanation of graphs see legend 233 to Figure 3.

As in Figures 3a, b, starting at 12 o’clock and proceeding clockwise, all the visits paid by each of the infants to these items of furniture are plotted in their order of performance, including the relative start and end times, and the extent of engagement exhibited in each of the visits. Generally speaking, the TD infants’ engagement agendas were much more eventful and dynamic than those of the NTD infants (evidenced by the number and variety of colored sections tiling the circles: multiple sharp spikes that touch the center in TDs (Almog, Dan, and Alon), versus a few extended sections of only a few colors that do not reach the center at all in NTDs (Adva, Dean, Evyater and Yuri). i) The TD infants visited multiple items of furniture, paying multiple visits to each; whereas the NTD infants visited the furniture much less frequently. ii) As evidenced by the extent of colored areas near the circles’ centers, the TD infants approached and often made contact with the furniture, whereas the NTD infants tended to approach the furniture less closely and engaged with it less deeply. iii) The large gray empty spaces in the NTD engagement graphs disclose the tendency of the NTD infants to sometimes adhere to places that were distant from the furniture, either engaging in stereotypies or engaging with a toy, or perhaps attending to themselves rather than to the environment. Adva was exceptional in the NTD group in visiting mother at regular intervals, albeit only for four visits. Two infants, Alexey and Shuval, paid multiple visits to the doorway leading out of the room, trying to open it, perhaps intending to leave. The engagement graphs also highlight significant within-group differences: e.g., while Almog alternated between visits to mother and single visits to an object across the session, Alon visited several items of furniture between successive visits to mother. The TDs took longer rests with mother, whereas the NTDs took longer rests near furniture.

### The infants’ management of distance from mother

We parsed the infant’s path into excursions by using the zero crossing of the infant’s path with the horizontal line marking the boundary occupied by mother’s customized place (see *Methods*). Thereby, touching, or crossing the line on the way down or on the way up defines a visit to mother. A segment of the path located above the horizontal line and bounded by two zero crossings defines an excursion. As demonstrated in Figure 5a, the most noticeable feature of the TDs exploratory path is its partitioning into excursions. Four of the TD infants (Alon, Almog, Dan, and Yoram) started the session with short duration excursions (marked by sharp peaks) and then proceeded to excursions involving extended lingering episodes (marked by flat-topped peaks). All (except Alexey) ended the session by cuddling on mother’s lap.

**Figure 5 a.**
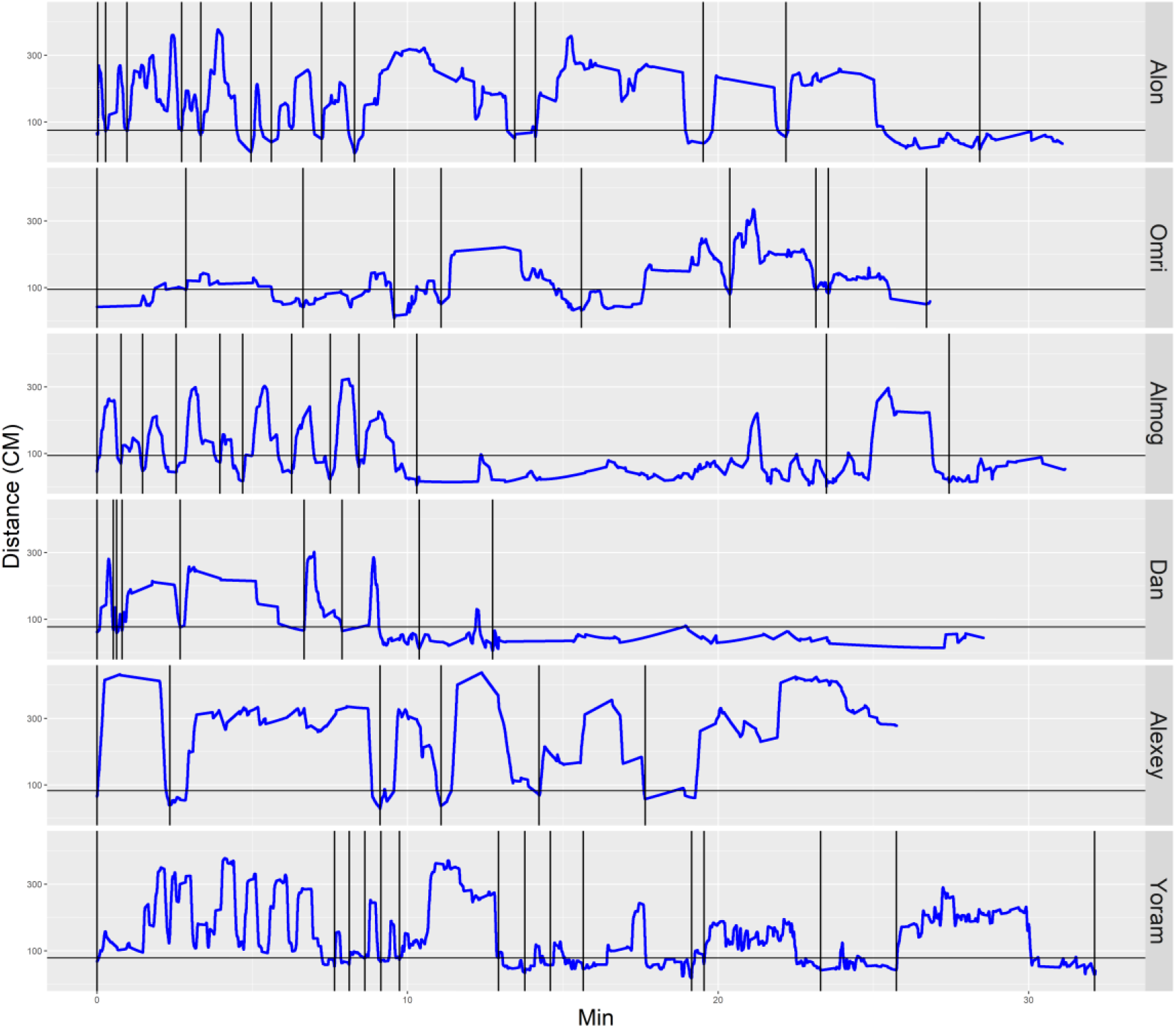
A plot of the TD infants’ management of distance from mother highlights the 289 multiple, repetitive outbound-inbound motion, the extent of engagement with mother (sections 290 below the horizontal line), and a tendency to start the session with sharp peaks (short durations 291 of staying at the far end of excursions) and continue with flat-topped peaks (extended durations 292 of staying at the far end of excursions; Alon, Almog, Dan, and Yoram). Alexey, exhibiting 293 several features of the NTD infants, is an exception, starting the session with an extended flat-294 topped peak (see *Methods*). Blue line plots distance from mother, black horizontal line marks 295 mother’s customized boundary, and black vertical lines segment the plot into excursions.

In contrast, the NTD infants (Fig. 5b) performed very few if any excursions from mother (the difference is 8.5 Wilcoxon rank test p-value 0.005): Adva performed four excursions ending up on mother’s lap; Dean did not visit mother for most of the session, climbing on her lap for nursing only during the last part of the session; Evyatar started the session with two excursions, but then avoided mother until the end of the session when he climbed on her to nurse; Tom established a place at a distance, vis-à-vis mother, which he visited several times, sitting there, facing mother, and performing stereotypic behavior without approaching her – as though there was an invisible glass wall between them; Yuri started the session with three incipient excursions (not captured by the algorithm) and then avoided mother for the entire session, chasing a ball, and ending the session by lying on his belly and crying in the middle of the room; and Shuval remained at a distance from mother for half of the session, visited mother only twice during the second half of the session, and spent a lot of time near the exit doorway, ending the session by crying by the doorway.

**Figure 5 b.**
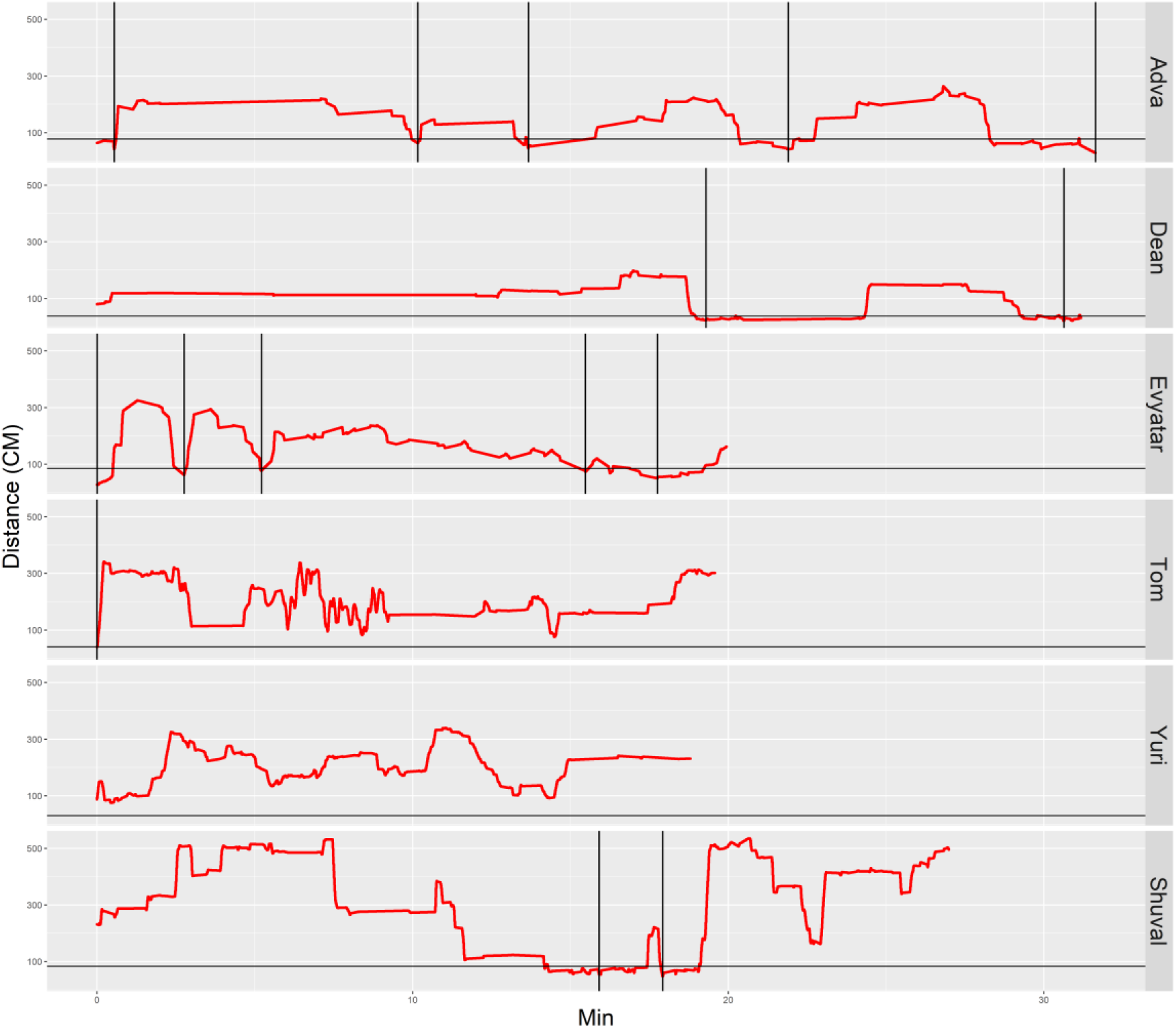
A plot of the NTD infants’ management of distance from mother highlights few if 298 any visits to mother, shallow engagement with the mother (hardly any sections below the 299 horizontal line), and a tendency to start the session with flat-topped peaks (long durations of 300 staying at a specific distance, indicating long staying in place episodes). Only three of the 301 infants end the session on mother’s lap. Red line plots distance from mother, black horizontal 302 line marks mother’s customized boundary, and black vertical lines segment the plot into 303 excursions.

The NTD plots are characterized by long straight lines that maintain a relatively fixed distance in reference to the mother’s location line, indicating that these infants are slow (See *Endpoints Summaries* – progression speed), walking away from mother and staying away for long durations whereas the TDs bounce back and forth across the y-axis, being more dynamic (See *Endpoints Summaries* –speed outside of mother’s vicinity). It should also be noted that the maximal distance from mother is much higher in the TDs, except for Shuval whose mother is located, unlike all the other mothers, at the opposite end of the room (see Fig. 2b).

### Physical contact with mother

Although visits to mother do not necessarily imply physical contact, a major difference between the TD and NTD infants was the amount of physical contact they established with their mother (See *Endpoints Summaries* – contact episodes and % of contact time). All the TD infants except for Alexey ended the session with a relatively long contact episode. In the NTD group only two infants ended the session that way. The other four infants ended the session by staying in place away from mother. It should be noted that some of the short physical contact episodes were initiated by the mothers, who leaned forward and established physical contact with their nearby passing infant.

### Excursions versus one un-partitioned path

The animations presented below demonstrate the partitioning of the path into excursions in two selected TD infants.

Animation: <u>Partitioning of Almog’s path into excursions</u> (TD infant)

Animation: <u>Partitioning of Alon’s path into excursions</u> (TD infant)

Animations of similar partitioning of the paths of all the other TD infants are presented in the supplementary material (S. videos 1-6). The TD infants exhibit a highly mother-centered organization involving excursions with clear outbound and then clear inbound portions.

The animations of the paths of two selected NTD infants presented below illustrate how the NTD paths tend to avoid mother’s place.

Animation: <u>Partitioning of Yuri’s path into excursions</u> (NTD infant)

Animation: <u>Partitioning of Tom’s path into excursions</u> (NTD infant)

Animations of similar, half-hour, un-partitioned paths of all the other NTD infants are presented in the Supplementary Material (S. videos 7-12). The NTD infants tend to avoid visits to mother.

## Discussion

### Classical infant-mother studies

: Most classical researchers of human infant mother-related exploration prioritize the analysis of its functional aspects, engaging with psychotherapy (Stern 2009), psychoanalytic theory (Mahler et al. 1975), or the study of an infant’s emotional life (Bowlby 2012), currently also in correlation to neural maturation (Schore 2015). This includes, for example, the study of attachment (Ainsworth & Bowlby 1991;Solomon & George 1999;Bowlby 2012), detachment (Rheingold & Eckerman 1970), separation-individuation in the infant’s intrapsychic life (Mahler et al. 1975), sense of infant’s self (Stern 2009), and inter-subjectivity (Trevarthen 1979). As such, these studies typically attend to multi-dimensional categorical prototypes involving expert evaluation, such as secure or insecure relationships (Solomon & George 1999), as well as to key episodes disclosing emotion and attention (Stern 2009). Using the researcher’s own free-floating attention "the psychoanalytic eye lets itself be led wherever the actual phenomenological sequences lead" (Mahler et al. 1975), ignoring spatiotemporal continuity. Whether reporting selectively or scoring in real time or from video, the classical researcher’s attention is typically attracted to the mother’s and infant’s face and hands, describing head movement, gaze, facial expression episodes, crying, vocalizing, and manual manipulation of objects (e.g., (Stern 2009;Mahler et al. 1975)), leaving the study of the structure of whole body movement, which carries along the head and the hands across the environment, to studies of locomotion (e.g., (Soska et al. 2010)) and exploration, who do not necessarily focus on the mother-infant interaction (e.g., (Gibson 1988;Kretch et al. 2014;Kretch & Adolph 2017;Adolph & Berger 2007)).

### “Structure-first” versus “function-first” paradigms

: Developmental psychologists aim at high level functions like intimacy and enduring emotional bonds. They would caution that even a temporary disregard of the infant’s attachment type may distort the interpretation of exploratory behavior (Cooper et al. 2011;Ainsworth et al. 2015). They might even require that a study of the strange situation would precede the study of the mother-infant situation, based on the claim that one does not know a phenotype unless one challenges it. From their vantage point, ignoring the nature of the emotional bond between the infant and its mother may distort the interpretation of the infant’s behavior.

At the other extreme, a comparative anatomist of behavior searching for the ancient phylogenetic origins of exploratory behavior would prioritize structure (Raff 2012;Gomez-Marin et al. 2016), and view the mother-infant situation as a basic, natural situation requiring structural characterization and understanding before rushing into the application of experimental perturbations. In this view, analysis of the basic situation across the phylogenetic scale in reference to origin-related exploration (Golani & Benjamini 2018) would be fundamental and primary. From this vantage point, even the presence of mother would, in a way, be dispensable had there been an alternative way of studying the infant’s exploration in the room by using an alternative attractor that would serve as a meaningful yet ethical origin and reference for the infant. From this perspective, challenging a phenotype before even isolating the perceptual quantities (order parameters) managed by it (Golani 1981;Powers 1973) would be senseless.

The fact that a controversy between structure-first and function-first paradigms prevails in the study of biological phenomena for almost three centuries (Appel 1987) implies that both hold a grain of truth. They can be viewed as complementary and/or useful depending on one’s aims. There is no a priori objective way of choosing between the two paradigms, except for by comparing their respective predictive powers. But whereas the first paradigm has been extensively used in the study of mother-infant interaction, the use of the second paradigm in a human context is too young to be evaluated, constituting only a first step in this type of structural comparison. Still, the structure-first paradigm deserves a chance, having proven itself in the study of animal origin-related exploration (Golani & Benjamini 2018), and providing a common framework for the study of organisms as varied as fruit flies, mice, primates and human infants.

### Current study

Complementarily, here we studied human infant mother-related exploration in a phylogenetic perspective, prioritizing the analysis of its structure (Amundson 2005), but in reference to mother. To accomplish this aim we focused on the scale of moment-by-moment (*actualgenese*; actual-genesis), using reports on close and remote taxonomic groups’ origin-related behavior as background material. Given our aim, a main requirement we fulfilled was to represent behavior in a way that would be useful in comparisons across taxonomic groups of, for example arthropods and mammals, including primates. The plots of the paths traced by the infants fulfilled this requirement. Using improved tracking technology, it should, however, be enhanced in future studies with concurrent separate representation of progression, of trunk orientation, and of the relations and changes of relation among all the moving parts of the kinematic linkage (Eshkol & Harries 1998;Golani 2012) (clearly also including hands and head movement, facial expressions, gaze and vocalizations, as well as a continuous record of mother’s visual attention, a highly important parameter (Sorce & Emde 1981). not examined in the current study). The highway methodology to deciphering the lived experience associated with the symptoms of sensory and motor differences (Donnellan et al. 2013) exhibited by the two respective groups of infants is a comprehensive Movement Notation analysis of the behavior (Eshkol 1958;Eshkol & Harries 1998), including the analysis of stereotypy (Golani et al. 1999).

Having analyzed the infants’ path in the environment we then isolated what appear to be candidate natural particulate processes (Wagner 1996;Golani et al. 1979) that make up the path in arthropods (Cohen et al. 2015) and vertebrates (Drai et al. 2000;Fonio et al. 2009;Golani & Benjamini 2018), constituting the elementary morphological characters of origin-related exploration (Gomez-Marin et al. 2015): progression segments and staying in place (lingering) episodes (Drai & Golani 2001), excursions (Eilam & Golani 1989;Tchernichovski et al. 1998;Benjamini et al. 2011;Drai & Golani 2001), and natural origins in reference to which the excursions are performed (Golani 2012) (see *Excursions versus one un-partitioned path*). This organized structure (Golani & Benjamini 2018) is used both in the same and in different organisms for different functions, such as management of novelty (Gordon et al. 2014) and of arousal (Fonio et al. 2009), socialization (Hinde & Simpson 1975) foraging (Woodgate et al. 2016), as well as the functions ascribed to human infants’ behavior (attachment, etc.). One major side benefit of this type of representation and analysis is that it is inherently translational, presumably consisting of the same cross-phyletic behavior (Golani & Benjamini 2018).

### Infant’s management of attention and of perception in reference to a stationary mother

We asked the seated mother to let her infant slide down from her lap and for her to remain in place across the half hour session, while we obtained a continuous track of the infant’s behavior on the path scale. Leaving the infants to their own devices had a profound effect on their behavior. Finding themselves in a novel, relatively pleasant environment, without being continuously bombarded by social stimuli, yet under the relatively silent visual attention of mother (only one or two of the mothers sporadically sank into reflection), the infants were entrusted with the full management, at their own pace and for an extended period of time, of their own location, distance, opposition (Yaniv & Golani 1987;Eshkol 1958) contact, and extent of engagement with mother, furniture, and toys, disclosing to the observer through their behavior the endogenous constraints that shaped their attention, perception, and engagement with the physical and parental environment. By mildly reducing the mother’s retrieval response in a safe environment we uncovered commonalities as well as an unsuspected disparity between the TD and the NTD infants.

### A computational analysis of the number of visits to the most preferred places in TD versus NTD infants

Making no prior assumptions regarding mother’s significance we first established each infant’s most preferred place in the environment, as reflected in the peak amount of cumulative time spent in it. To ascertain that the place was indeed preferred persistently across the whole session we also required that dwell-time would be incrementally accumulated in that place through a sequence of visits that would be more-or-less evenly spread across the session. Places presenting these features were located in mother’s proximity in five of the six TD infants but in none of the NTD infants (figs. 2a, b). Having established the preference for mother’s proximity in the TD infants, we then plotted in the order of performance the time of start, the duration, the time of end, and the extent of each of the infant entries into mother’s proximity (Figs. 3, 4). In addition, we calculated the number of visits to mother by establishing a customized (Benjamini et al. 2010) boundary for each infant-mother session (see "segmentation to excursions" in *Methods* and Fig. 5). Finally, we found that the TD infants paid a higher number of visits to mother at all distances regardless of where we had established a boundary (fig. S1). Most of the TDs’ mothers were thus visited persistently across the session, whereas most of the NTDs’ mothers were visited rarely and irregularly if at all (see boxplot summaries Fig. 7).

**Figure 7.**
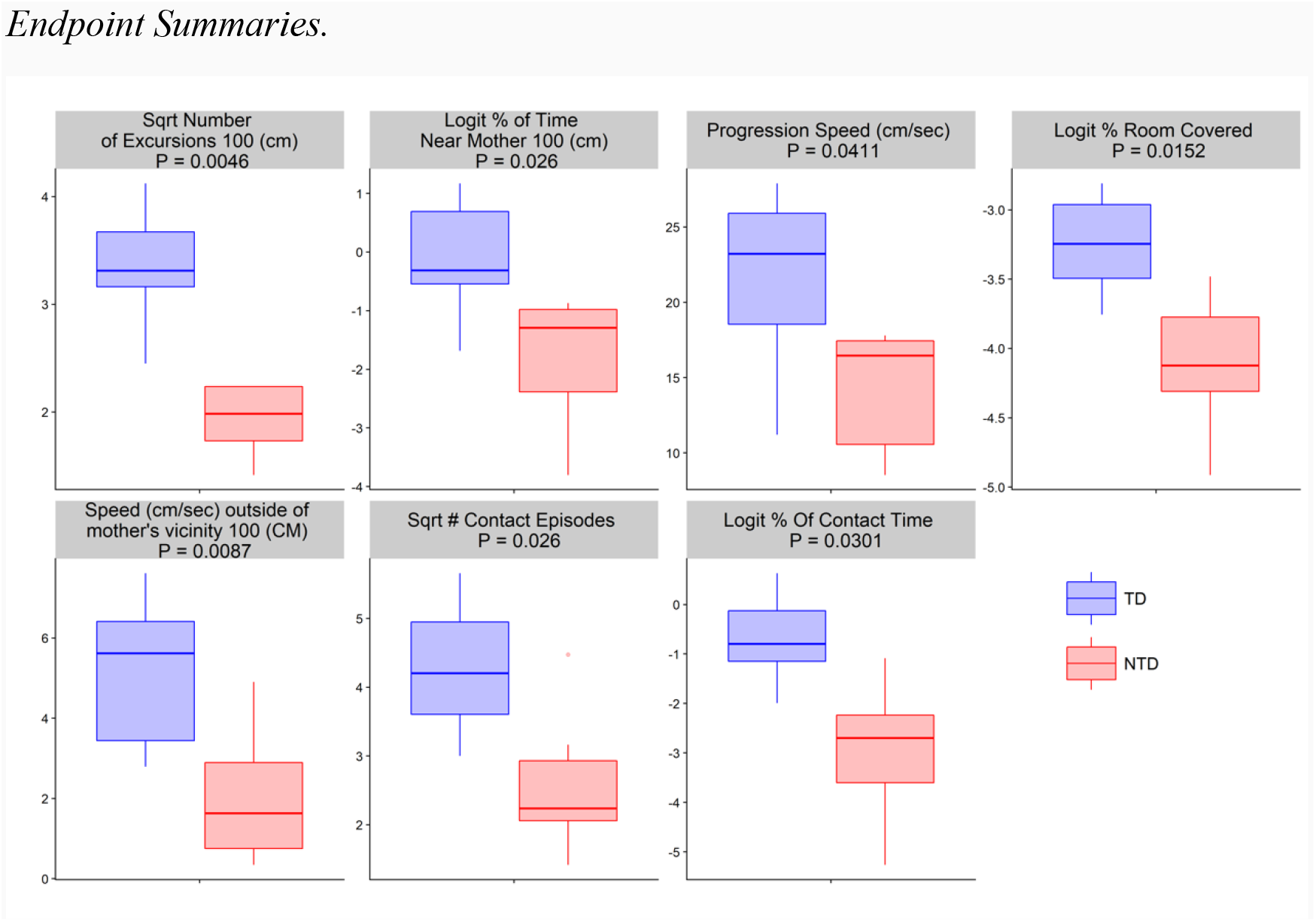
Boxplot summaries demonstrate significant differences between the TD and NTD 332 infants’ behavioral endpoints. All comparisons were conducted using Wilcoxon rank test 333 (requires no assumption regarding the underlying data distribution). Across all endpoints the 334 differences between the TD & NTD infants are significant (at significance level 0.05) after 335 correction for multiple comparisons (Benjamini & Hochberg 1995). The comparison between 336 the number of excursions and % dwell time is dependent on the radius that defines the mother’s 337 location. The difference remains robust to changes in that radius (see Fig. S1 and associated 338 text in *Supplementary Information*).

### Partitioning the flow into excursions

Visits to mother were next used to partition the overall path into excursions that started and ended in mother’s vicinity (vertical bars in Fig. 5a, b). The segmentation of the path into excursions reveals that the TD infants performed a median of 12±5 excursions per session whereas the NTDs perform 3±3; three of the NTD infants did not perform any excursions at all. Furthermore, the TDs come to extensive grips with mother, not only by physically touching her during visits (fig. 6) but also by climbing and thus deeply engaging her (Figs. 3, 7). Some of the TD infants, like Almog, performed relatively simple excursions composed of clear, relatively monotonical outbound and monotonical inbound portions in reference to mother: their exploratory path is tightly centered on mother. Others, like Alon perform both simple and complex excursions that include several non-monotonical back-and-forth shuttles (Fonio et al. 2009) on the inbound portion of the excursion. Nonetheless all the TDs’ exploratory paths were tightly centered on mother.

**Figure 6.**
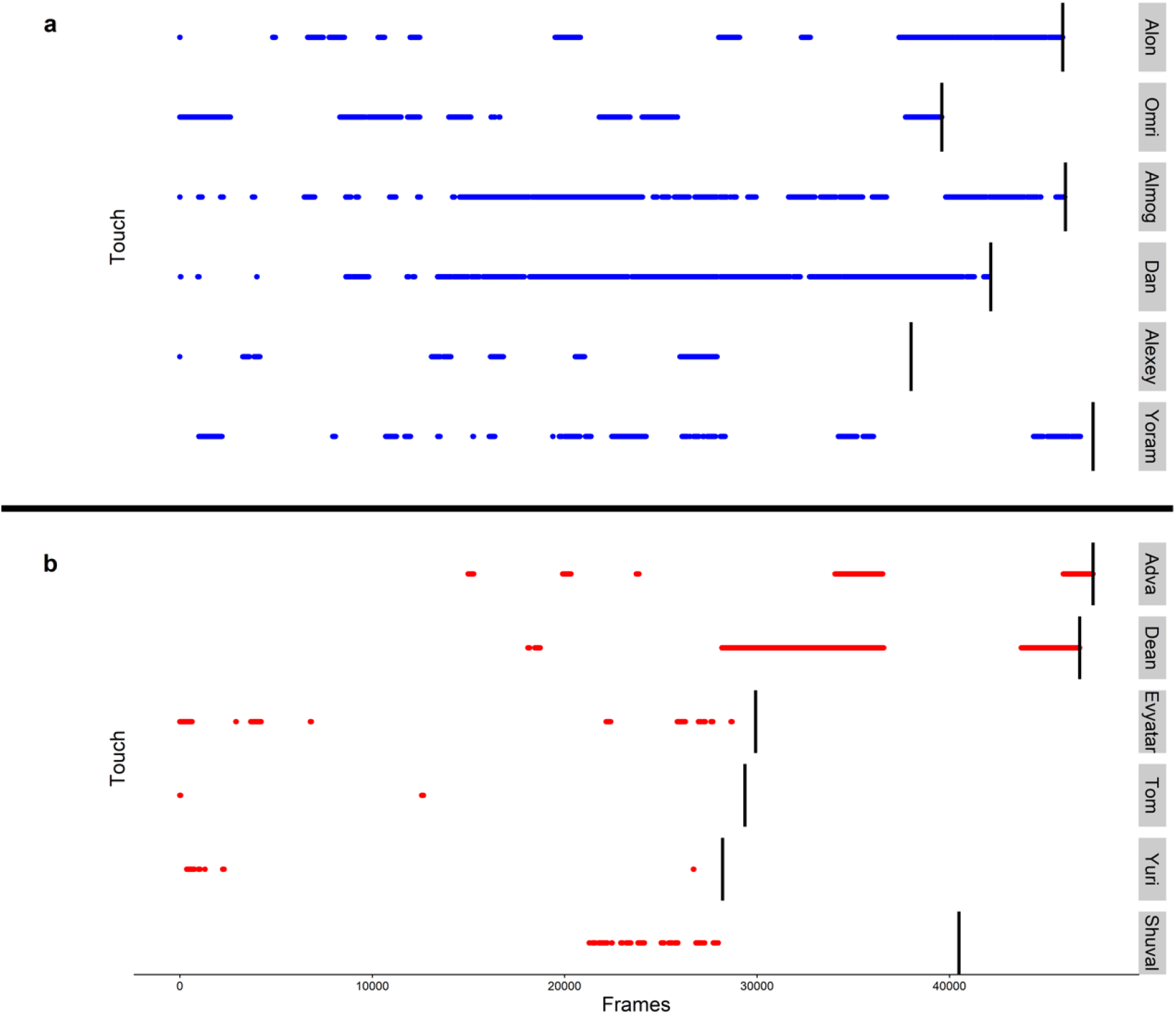
The duration and frequency of episodes involving physical contact with the mother 326 were high in the TD infants and low or almost absent in the NTD infants. The vertical black 327 lines represent the end of the respective infant’s session.

### The importance of excursions in infant-mother relationship

In TD human infants the excursions are both the units of exploration of the environment and the units of discourse with mother. They are repeated sequences of alternating interactive units of experience with mother and with the world, encompassing both the world and mother in the same excursion. Mother experience and world experience come in pairs. The excursion is a manner of interaction between infant and mother, a “schema-of-being-with-mother" (Stern 2009), managed mostly by the infant, who "makes love" with the environment and "rushes to tell mother", much like a playing together or a sharing-of-feeding episode. It is a sequence that takes on "a regular, almost canonical form that can become an internalized model used to evaluate current experiences” (Stern 2009). The "checking back pattern" is the most important fairly regular sign of beginning somato-psychic differentiation, according to Mahler, who considers it to be the most important normal pattern of cognitive and emotional development. Its central role is expected considering that for most of human and primate history infants were probably carried by their mothers, using mother as an origin during forays into the environment, unlike the infants of other mammalian orders that either follow mother or are nested, or being cached (Jones 1972). Examination of mother-related exploration from a phylogenetic perspective reveals that for human infants in specific contexts, origin, mother, and homebase are one.

### The ontogeny of primate infant mother-related excursions

During the first postnatal period, mothers of monkeys, baboons, and chimpanzees carry their offspring continuously, until such time as the infant descends from mother, making increasingly longer excursions away from and back to her (Altmann 2001;Plooij 1984;Berman 1980). In chimpanzees, starting from the 8^th^ month and on, the infants make short excursions remaining within arm’s reach from their mothers, and then, across months, gradually increase the frequency and maximal distance of excursions to the point that the mothers might be out of sight of the infant for extended durations (Plooij 1984). A similar growth in the frequency and extent of excursions has been documented for a variety of monkeys kept in captivity (van Lawick-Goodall 1968;Nicolson 1982;Pusey 1978;Kaufmann 1966;Hinde & Simpson 1975;Rheingold & Eckerman 1970). Even in controversial rearing-in-isolation experiments, in which the monkeys were reared in the presence of artificial surrogate mothers (made of wire), the infants used the surrogate as their "base of operations", moving away to examine furniture in the environment and then returning to base (Harlow & Zimmermann 1959).

In all the mother-infant interactions in primates, both partners initiate departures and approaches, contributing to the management of distance and contact between them (Plooij 1984;Hinde & Simpson 1975). In rhesus monkeys, the relative proportion of infant initiated departures and approaches increases with age (Hinde & Spencer-Booth 1967). One way by which infant chimpanzees manage the distance with mother is by whimpering and inducing retrieval (Plooij 1984). The human mother appears to retrieve the infant during the pre-walking stage less often than many other mammals do, being present in environments in which restraining and retrieving are less necessary. One third of the inbound trips in 10-month old human infant excursions did not end with actual physical contact: "to see seemed sufficient (Rheingold & Eckerman 1970). Human infants, unlike many other mammals, venture to explore away from their mothers from the moment that any mode of locomotion becomes possible (Rheingold & Eckerman 1970); they do not wait until they can creep or walk efficiently.

### The phylogenetic status of human infant mother-related exploration

: The claim that mother-related exploration is homological across the primates is supported by the distinct taxonomic distribution of the carrying plus using mother as origin plus shared alternation between progression and lingering, plus excursions, all vis-à-vis the distinctly different taxonomic distribution of following, nesting, and caching (Jones 1972). The claim that this behavior is homological to origin-related exploration in fruit flies, crustaceans, rodents, and fish is supported by the shared connectivity (Wagner 1996;Saint-Hilaire 1822) characterizing this behavior across phyla. The wide range of functional contexts in which the same structure unfolds is an essential characteristic of evolutionarily conserved characters: in the context of developmental evolutionary biology origin-related behavior is a character identity (Wagner 1996) supporting different functions. Establishing the evolutionarily-conserved status of human infant mother-related exploration and of its constituents would provide a step forward in the search for a homeomorphic organization that might mediate this plan’s architecture in the brain.

The paucity or absence of excursions in the NTD infants is remarkable in view of the universality of mother-related exploration in primates and the sharing of origin-related behavior in vertebrates and arthropods. In the absence of a preferred stable place, the NTD infants’ paths could not be segmented into excursions: they were punctuated like those of the TDs paths by lingering episodes, but these episodes tended to be situated away from mother. As soon as the session started, both TDs and NTDs slid down from mother’s lap and crawled away, but while the TDs tended to immediately progress back and forth in reference to mother, lingering briefly in several locations along the excursion’s path, the NTDs performed an extended staying-in-place episode as soon as they reached a piece of furniture or a toy (Adva, Tom, Evyater and Shuval), or as they stopped in an empty space away from walls (Dean), and tended to stay in place at that location for extended durations (Fig. 5b). Such extended staying-in-place episodes are also performed by TD infants, but typically much later in the session, only after performing a sequence of short staying-in-place episodes (Fig. 5a Alon, Almog, Dan and Yoram). The paucity or absence of visits to mother in the NTDs does not necessarily imply ignoring mother: a preliminary analysis of infant gaze suggests that these infants gazed at mother from across the room at least as often as did the TD infants. The "glass wall" surrounding the NTD mothers (e.g., Tom and <u>Yuri</u>) appears to disclose a mixture of active avoidance of proximity, including partial or full evasion of physical contact with mother (fig. 6), and an enhanced focused attention on objects.

Left to their own devices under the extended *laissez-faire* attention of mother, the operational worlds of the NTD infants unfolded as a single, undividable, attentive yet haphazardly oriented (exhibiting no reference to an origin) slab of behavior. To the best of our knowledge no vertebrate or arthropod has been reported to date to exhibit such behavior, nor have we been able to identify an animal model, be it a fruit fly, a rodent, or a monkey, that performs active investigation of the environment without referring to an origin, or while being relatively free of the attraction of an origin. The search image portrayed by the present study for such model thus shapes up as an animal capable of switching origin-related behavior off.

The intriguing findings by the current study could perhaps lead us to associate the described behavior of the NTD infants with early signs of autism, as these infants were referred to the Mifne Center for Early Treatment of Autism for assessment and/or treatment. However, such a conclusion is beyond the capacity and confines of this study. In order to make such an association it would be necessary to replicate (Kafkafi et al. 2017) the study with large groups of TD and NTD infants, concurrently performing follow-up developmental tests at older ages, and screening for developmental disorders, including Autism Spectrum Disorders (ASD).

## Experimental design & methods

Subjects were twelve 8-12 month-old pre-walking infants recruited with their respective mothers to participate in a videotaped half hour session conducted at the Mifne Center. All infants were documented at the stage of stable crawling. Six of the subjects wer Typically Developing (TD) infants whose parents had volunteered to participate in the experiment. None of the TD infant parents had raised any concerns regarding developmental problems. The other six subjects were Non-Typically Developing (NTD) infants who had applied to the Mifne center for developmental assessment and/or treatment, following referrals by child neurologists (three infants) and expert clinician (three infants). The parents were asked to sign a form designed by us and approved by the Ethics Committee of Tel Aviv University. The NTD infants underwent external expert developmental assessments, including the verification of inclusion and exclusion, detailed family history, perinatal, medical, and developmental history, and physical and neurological checkups. For each trial the mother entered the trial room carrying her infant sat on a mattress in the periphery of the room, and then seated the infant next to her. The mother had been requested to remain seated within the area of the mattress and allow the infant to act freely for a 30-minute session. Not all sessions lasted for half an hour, with some being stopped if the infant appeared to be distressed (see Figs. 5, 6). Two video cameras, one capturing a static view of the whole room including the infant and the mother (fig. 1), and the other zooming in on the infant and following it, were used across the session. The static camera data were transformed to a top view (see below). Data were prepared for segmentation (Hen et al. 2004;Drai et al. 2000;Drai & Golani 2001) and were subjected to analysis by SEE, a publicly available Software for the Exploration o Exploration developed and elaborated by us over the course of many years (www.tau.ac.il/∼ilan99/see/) (Drai & Golani 2001).

### Room

The infants were tracked in a medium-sized, 3.65m by 5.45m room.

### Tracking

The tracking of the infant was done manually using a specifically dedicated program in Matlab. The infant’s location was tracked every ~15 frames; missing coordinates were completed using linear interpolation.

We resorted to the use of manual tracking after exhausting other possibilities: different tracking algorithms had failed due to bad image quality, multiple object moving and the fact that an infant is a large object, so even when tracking was successful, there was jitter across the infant’s body.

### Transformation

The room was videotaped from a high side view camera angle requiring a projective transformation to an exact overhead view that would replace the 2D coordinates in the video image (Fig. 1, left) to 2D coordinates in the world (Fig. 1, right). A projective transformation is an image-processing technique that finds the linear transformation from one coordinates basis to another. It was implemented in Matlab, by choosing 4-5 points (whose coordinates were known for both bases) and finding the projection matrix.

### Cumulative dwell-time maps (heat maps)

The construction of the heat maps involved several steps:

1. Obtaining the original coordinate data from the Matlab tracking.
2. Smoothing the coordinates using the SEE program (https://www.tau.ac.il/∼ilan99/see/).
3. Dividing the room into a grid, in which each cell is a 1 (*CM*) ∗ 1 (*CM*), and calculating the time spent in each cell according to 2.
4. Smoothing the cells using 2d Gaussian smoother: calculating for each cell a new value according to the weighted average of the cell itself and its neighboring cells. The weights are given by the a 2d Gaussian distribution (σ^2^ = 14, *n* = 63) (fig. 8)
5. Finding local maxima of the smoothed coordinates and discarding the 96% of maxima’s with lower values.
6. Calculating the number of visits to each local maximum, based upon the two concentric circles (*r_in_* = 30, *r_out_* = 50) paradigm described below.

We used several distinct ways for visualizing and scoring the infants’ visits to places and for segmenting the path traced by them across the session. We calculated the number of visits to peak dwell-time places across the room using the two concentric circles algorithm; we used engagement plots to visualize all the infant’s entries into mother’s close proximity, including their extent and duration; and we established a customized boundary for each mother’s place in order to calculate the number of visits paid to mother and the number of excursions or round-trips performed from her into the environment.

### Using two concentric circles to calculate the number of visits to a place

The number of visits to a circumscribed area could be calculated by selecting a location, tracing a radius around it, and counting the number of times the infant entered the defined circular area. However, small vacillations of the infant across the border of the circular area without actually crawling away and returning would count as distinct, multiple visits. To avoid this outcome, we defined two concentric circles, both centered around the same location of interest. A visit started when the infant entered the inner (*r_in_*) circle and ended when it exited the outer circle (*r_out_*).

### Engagement Plots

The engagement plot exhibits the timing, duration, and extent of the infant’s being in gear with mother and with furniture and doors in the room. Each set of concentric circles exhibits the behavior of a specific infant. Starting at twelve o’clock and proceeding clockwise for the session’s duration, the arc traced on the circle’s circumference and the colored section of the circle, designate the time of start, the time of end, the duration in session percentage, and the extent of the entry into mother’s or any other large object’s close proximity (the infant seemingly casts a shadow on the peripheral area separated from the circle by its entry). The color of the polygon stands for the visited object, with mother being colored in black. Furniture items are designated by specific colors. For radius *R*, time *t* and for object *j* a point is drawn according to (*D*_*j,t*_ < *R*), *D*_*j,t*_ being the distance of the infant at time *t* from the center of object *j*. In order to compare multiple pieces of furniture of different sizes, the distance drawn on the plot are the distance of the infant from the object minus the radius of the object. See Figure S2.

### Segmentation to excursions

To partition the infant’s path across the session into excursions we needed to define for each mother the customized circumscribed place she occupied. The length of the radius tracing the boundary of mother’s place in reference to mother’s center influenced the number of visits paid by the infant to mother across the session: the smaller the radius the fewer the number of visits. To obtain a customized place around the location defining mother’s center, we progressively increased the radius circumscribing mother’s location, dynamically changing the number of visits paid to mother. The minimal radius yielding a relatively stable number of visits for a vector of protracted lengths of radii was selected as the radius defining mother’s place. We obtained the number of visits by using the two concentric circles method for each inner circle radius between 30 cm to 120 cm and an outer circle radius that was larger by 1.1 than the inner circle radius, then searched for the longest radii interval in which there was no change in the number of visits, and then chose the smallest radius of that interval.

## Data Availability

All data generated or analyzed during this study can be found at https://github.com/tfrostig/Infants-Analysis.

## Acknowledgments

The development of SEE software has received further funding from the European Research Council: ERC grant agreement PSARPS{294519}.

This work continues the pioneering study conducted by Dr. Hani Monk-Vitelson supervised by Prof. Yuval Portugali and Prof. Chaim Pick of Tel-Aviv University and two of the current co-authors. We thank Hagar Yulzari, Dr. Neri Kafkafi, and Dr. Hillel Dr. Braude for their useful comments on the manuscript and Shana Salomon for the acquisition of the physical contact data. Ms Naomi Paz edited and proofread the manuscript. We thank the Mifne Center’s personnel for their commitment, help, and enthusiasm throughout the study.

## Supplementary Information

### Quantitative examination of dwell time distribution in reference to mother

To quantify dwell-time, we plotted the average time spent across the session for the whole range of distances from mother’s center to a distance of up to 1.2m. The results reveal that the TD infants spent significantly more time than the NTD infants within the entire range marking mother’s vicinity. Only the exceptional TD infant Alexey (see *Methods*) shows the lower values characterizing the NTD infants; the lowest curve for both number of visits and dwell-time in the TD infants belongs to Alexey. Using permutations to test the difference between the two average curves is significant (p-value 0.029, the average difference between the curves is 0.19).

**Figure S1 a, b, c, d.**
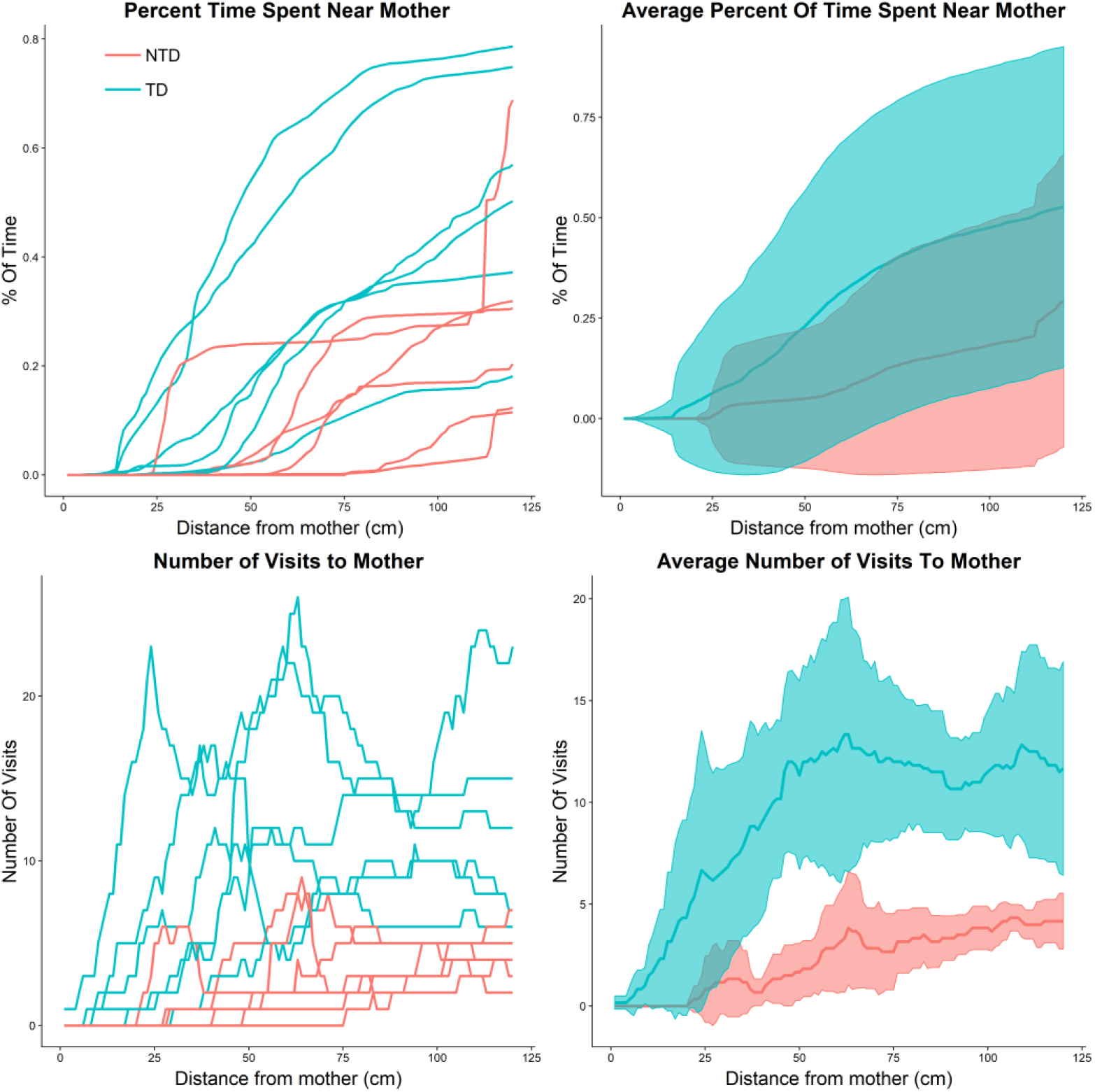
Percentage of time spent in mother’s vicinity. **c, d**: Number of visits to mother as a function of distance from mother. The CI for figures **b** and **d** was obtained using the normal approximation.

### Quantitative summary of number of visits-to-places distribution in reference to mother

Plotting the number of visits paid by each of the infants to the mother as we dynamically changed the inner radius of the two circles algorithm (the outer radius was kept at 1.1 of the inner radius) reveals that the number of visits to mother are higher for the TD compared to the NTD infants for the whole range of radii up to 1.2m. Since excursions are punctuated by visits to mother (see section *The infants’ management of distance from mother*), it also follows that for all considered distances from mother the TD infants exhibit more excursions than the NTD infants. Using permutations to test the difference between the average excursions curves of the TD and NTD is highly significant (p-value - 0.005, the average difference between the curves is 7.23 visits).

### Engagement Plots

**Figure S2.**
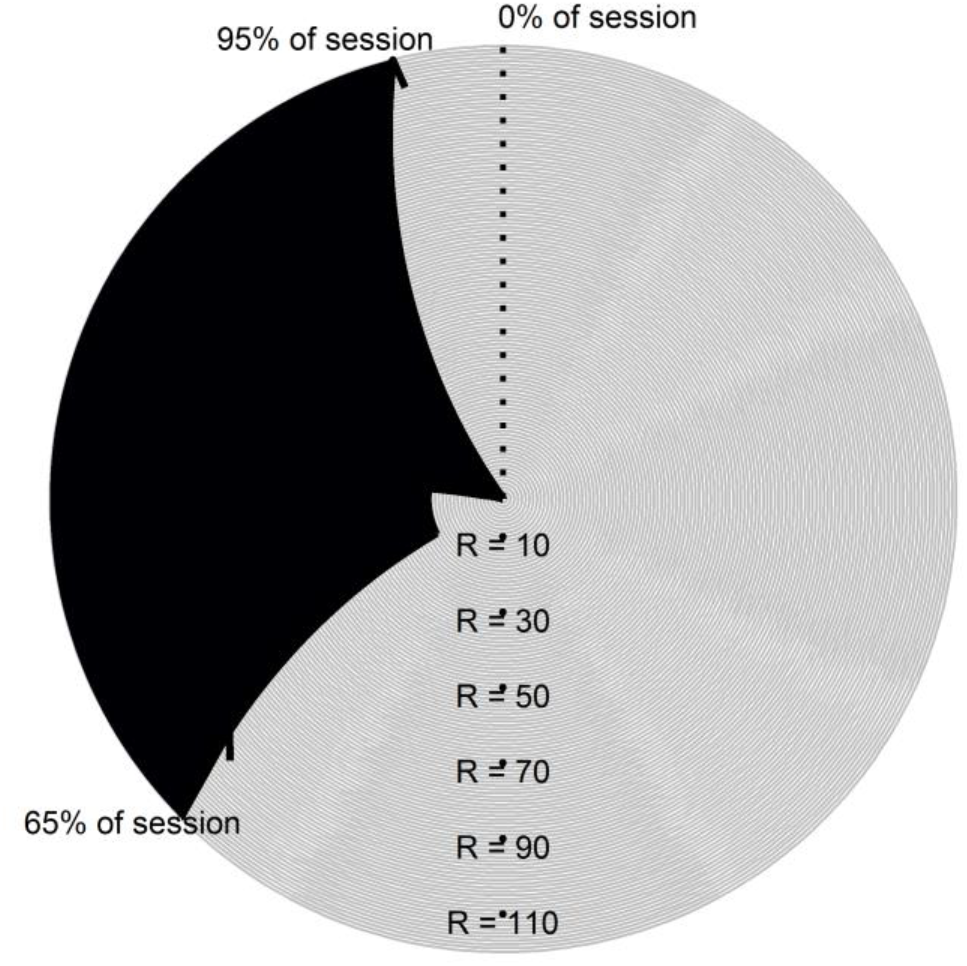
Illustration of an engagement plot including annotation. The black wedge represents the % of time the child spent near mother; the concentric circles radii represent the distance of the infant from mother; and the length of the arc at the respective radius represents the % of time spent at that distance (or greater).

### Videos

Animations of the infants’ sessions were created and can be viewed using the following links. Note that the infant sometimes appears to be beyond the boundary of the room, which is due to the tracking of the child and then transforming the image, so that when the infant is standing it appears that the tracker is beyond the boundary. The mother’s size changes from video to video, due to the way in which each mother sat (some were almost supine).

The infant’s center of mass in the current frame is represented by a black circle; movement in the current excursion is segmented into lingering segments; progression segments are represented by blue and red colors respectively, past excursions are shown in grey; the mother location is in green and the rest of the furniture items are in black.

### TD infants

<u>Video S1 - Alon animation of excursions</u>

<u>Video S2 - Omri animation of excursions</u>

<u>Video S3 - Almog animation of excursions</u>

<u>Video S4 - Dan animation of excursions</u>

<u>Video S5 - Alexey animation of excursions</u>

<u>Videos S6 - Yoram animation of excursions</u>

### NTD infants

<u>Video S7 - Adva animation of excursions</u>

<u>Video S8 - Dean animation of excursions</u>

<u>Video S9 - Evyatar animation of excursions</u>

<u>Video S10 - Tom animation of excursions</u>

<u>Video S11 - Yuri animation of excursions</u>

<u>Video S12 - Shuval animation of excursions</u>

### Statistical Testing

Tables for all tests conducted with additional information.

All tests are conducted with *n_x_* = 6 TD infants and *n_y_* = 6 NTD infants. SD is the pooled standard deviation 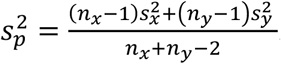.

The infants’ management of distance from mother

| Test – Wilcoxon rank-sum test | P-value | Means Difference<br>$\bar{x} - \bar{y}$ | Statistic (W) | SD<br>$s_p$ | Effect Size<br>$\frac{\text{Median}(x_i - y_j)}{s_p}$ |
| --- | --- | --- | --- | --- | --- |
| # of Excursions | 0.004922 | 8.5 | 36 | 2.783 | 3.233 |

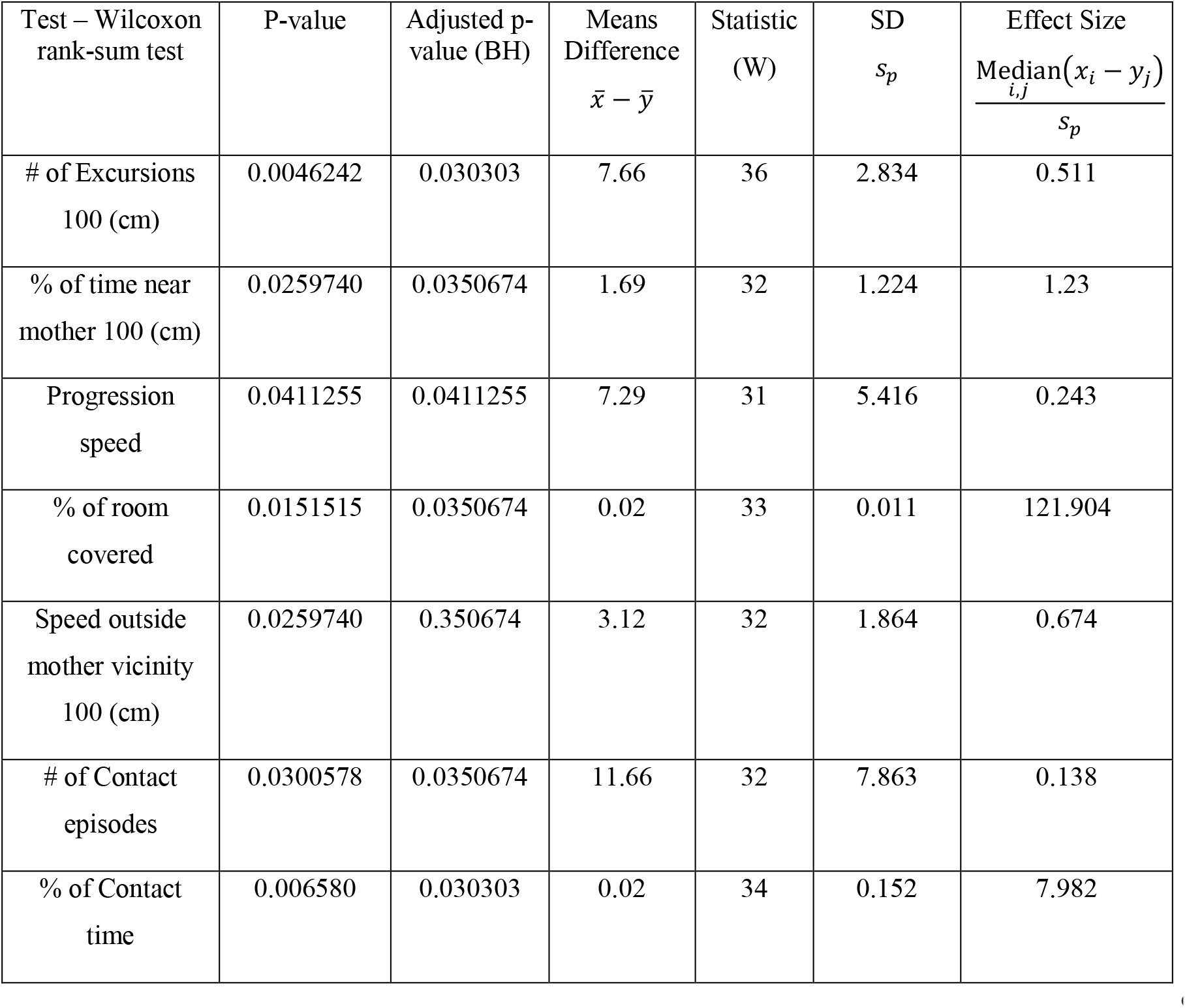
Endpoint Summaries.

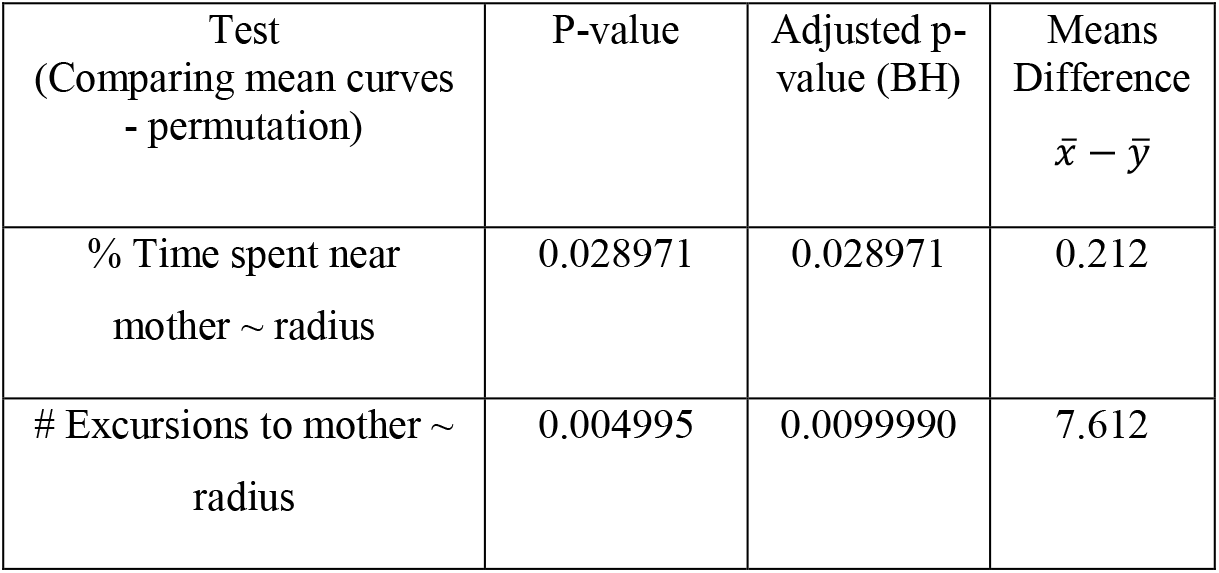
Quantitative examination of dwell time distribution in reference to mother and Quantitative summary of number of visits-to-places distribution in reference to mother

